# Predicting Cancer Drug Response Using a Recommender System

**DOI:** 10.1101/215327

**Authors:** Chayaporn Supahvilai, Denis Bertrand, Niranjan Nagarajan

## Abstract

**Motivation:** As we move towards an era of precision medicine, the ability to predict patient-specific drug responses in cancer based on molecular information such as gene expression data represents both an opportunity and a challenge. In particular, methods are needed that can accommodate the high-dimensionality of data to learn interpretable models capturing drug response mechanisms, as well as providing robust predictions across datasets.

**Results:** We propose a method based on ideas from “recommender systems” (CaDRReS) that predicts cancer drug responses for unseen cell-lines/patients based on learning projections for drugs and cell-lines into a latent “pharmacogenomic” space. Comparisons with other proposed approaches for this problem based on large public datasets (CCLE, GDSC) shows that CaDRReS provides consistently good models and robust predictions even across unseen patient-derived cell-line datasets. Analysis of the pharmacogenomic spaces inferred by CaDRReS also suggests that they can be used to understand drug mechanisms, identify cellular subtypes, and further characterize drug-pathway associations.

**Availability:** Source code and datasets are available at https://github.com/CSB5/CaDRReS

**Supplementary information:** Supplementary data are available online.

## 1 Introduction

Cancer is a genetic disease caused by the accumulation of mutations, ranging from point mutations to copy number variations and structural alterations. These, in turn, impact gene expression and ultimately contribute to the hallmarks of cancer, including uncontrolled cell proliferation and metastasis. Compared to commonly used cancer treatments such as chemotherapy or radiotherapy, targeted drugs can be better at killing tumor cells and/or have lesser toxicity to normal tissues (Begg *et al.,* 2011). However, not every patient responds to drug therapy in the same way, and molecular information such as mutation or gene expression data can inform us on which patients will respond to a drug. For example, KRAS mutations can be used as predictors of resistance to therapy with EGFR Inhibitors (Massarelli *et al.,* 2007), and targeting overexpressed Bcl-2, as observed in small-cell lung cancer, has been shown to provide therapeutic benefits (Gandhi *et al.,* 2011). These findings emphasize the need for using molecular information to predict drug response and thus personalize cancer therapy (Thangue and Kerr, 2011; Veer and Bernards, 2008).

As the number of patients/tumors with molecular data increases across cancer types, enabled particularly by large-scale studies such as TCGA and ICGC (Weinstein *et al.,* 2013; Zhang *et al.,* 2011), the identification of cancer driver genes has benefited greatly (Cerami *et al.,* 2012; Zhang *et al.,* 2011; Weinstein *et al.,* 2013; Bertrand *et al.,* 2017). However, these data sources typically lack drug response information and are therefore not suitable for identifying drug response biomarkers. On the other hand, drug screening on several panels of cancer cell lines has been conducted, for example, in the Cancer Cell Line Encyclopedia (CCLE) and the collaborative Genomics of Drug Sensitivity in Cancer (GDSC) projects (Barretina *et al.,* 2012; Iorio *et al.,* 2016). These cell line datasets allow us to utilize genomic features and apply mathematical and statistical approaches to decipher functional relationships and construct models that can predict patient-specific drug responses.

Several types of models have been proposed for predicting drug responses using genomic features (Azuaje, 2016; Costello et al., 2014; McLeod, 2013; Wheeler et al., 2012). The most widely used type is a drug-specific model, which is independently trained for each drug based on genetic and drug response information from cell lines tested with each drug individually. Some of the methods that fall in this category include, a linear regression model using baseline gene expression (Barretina *et al.,* 2012; Iorio *et al.,* 2016; Geeleher *et al.,* 2014) or based on a combination of gene expression and other genomic information such as copy number alterations and DNA methylation (Ding *et al.,* 2016; Chen and Sun, 2016), nonlinear models such as neural networks, random forests, support vector machines and kernel regression based on multiple types of genomic information (Cortés-Ciriano *et al.*, 2016; Dong *et al.,* 2015; Gupta *et al.*, 2016), and a neural network model that also incorporates drug property information (Menden *et al.,* 2013).

Drug-specific models are typically limited by the number of cell lines that have been tested with a given drug. To increase the number of data points and obtaining more robust and general models for drug response, a Bayesian multitask multiple kernel learning (BMTMKL) approach was proposed and exhibited the best performance in the DREAM challenge for drug response prediction (Costello *et al.,* 2014). This work highlighted the importance of sharing information across drugs in improving the accuracy of drug response prediction.

Multitask learning assigns all drugs equal importance in response prediction for a given drug, but it is likely more meaningful to construct a model that prioritizes information from similar drugs, as is possible using collaborative filtering techniques. In the area of recommender systems, collaborative filtering is a framework to analyze relationships between users (cell-lines/patients) and dependencies among items (drugs) to identify new user-item associations (patient-specific drug response) (Koren *et al.,* 2009). The two major classes of collaborative filtering techniques are (i) neighborhood methods, which predict the user-item association based on predefined user-user and item-item similarities, and (ii) latent factor models, which use matrix factorization to identify a latent space that captures user-item associations. Matrix factorization techniques, in particular, have shown promising results in the Netflix Prize, a competition for collaborative filtering methods to predict user ratings for movies based on a rating history (Bennett and Lanning, 2007).

Collaborative filtering techniques have also been used for predicting patient-specific drug responses in a few studies. Based on a neighborhood approach, Sheng *et al* (Sheng *et al*., 2015) defined drug-specific cell line similarity and drug structural similarity, and then predicted unobserved drug responses by calculating a weighted average of observed drug responses according to both drug and cell line similarity. This model is purely based on the assumption that the predefined similarities can explain drug responses, but it did not take into account observed drug response information to define drug similarity. In contrast, using the latent factor approach, Khan *et al* (Khan *et al.*, 2016) constructed component-wise kernelized Bayesian matrix factorization (cwKBMF) models to predict unobserved drug responses based on multiple cell line kernels and observed drug response data. Khan *et al* showed that cwKBMF can identify drug-pathway associations and outperformed BMTMKL (Costello *et al*., 2014) in drug response prediction. However, a common limitation of both models is a need for normalization of drug response data, with this preprocessing step leading to a loss of information on relative ranking of drugs within each cell line. Overall, the availability of limited training data, with a small number of cell lines tested with each drug, represents a major challenge for learning robust models that provide meaningful predictions in new datasets. Additionally, the interpretability of models and their use to obtain biological insights has not been extensively explored in the field.

To address these limitations and to develop more robust models based on information sharing across multiple drugs, we developed the CaDRReS (for **Ca**ncer **D**rug **R**esponse prediction using a **Re**commender **S**ystem) framework. CaDRReS maps drugs and cell lines into a latent “pharmacogenomic” space to predict drug responses for specific unseen cell lines and patients. Our benchmarking analysis using publicly available datasets (CCLE, GDSC) suggests that this allows CaDRReS to have notably better predictive performance and robustness than other existing methods. Comparisons on unseen patient-derived cell-line datasets also highlight CaDRReS’s robustness and ability to generalize across datasets, an important requirement for precision oncology applications. Additionally, we show that the unique pharmacogenomic space model inferred by CaDRReS lends itself well to biological interpretation, allowing us to (i) understand drug response mechanisms, (ii) identify cellular subtypes from drug response profiles, and (iii) characterize drug-pathway associations.

## 2 Methods

### 2.1 Datasets and Data Preprocessing

Drug-screening data for cancer cell lines was obtained from two large-scale studies, CCLE and GDSC, and all cell lines with baseline gene expression data were retained. A Bayesian sigmoid curve fitting approach was applied to raw intensity data at different drug dosages to recompute *IC*_50_ (minimal concentration that induces 50% cell death) values that were more comparable across datasets (see **Supplementary method 1, Supplementary figure 1** and **Supplementary tables 1-2** for details). Drugs with median *IC*_50_ less than 1 μM tend to be cytotoxic drugs with high toxicity across cell lines and were therefore excluded **(Supplementary figure 2).** Our final dataset contained 491 cell lines, 19 drugs, and 9,096 experiments from CCLE, and 983 cell lines, 223 drugs, and 179,633 experiments from GDSC, providing a large dataset for training and validation of our models. Additionally, an in-house dataset based on screening of 276 drugs (65 of which overlap with GDSC) on 8 head and neck cancer (HNC) patient-derived cell lines from 5 subjects was used (Chia *et al.,* 2017). Two of the cell lines were found to be not sensitive to any of the overlapping drugs (inhibition score <50 at 1 *μM*), while one was found to be sensitive to more than 25% of the overlapping drugs. Excluding these, 325 data points from 5 cell lines were used as an independent dataset to evaluate predictions from different models.

### 2.2 Cancer Drug Response prediction using a Recommender System

The first step in CaDRReS is to calculate cell line features based on gene expression information. To do this, we normalized baseline gene expression values for each gene by computing fold-changes compared to the median value across cell lines. For the next step, since the drug response experiments in GDSC and CCLE aim to measure cell death, 1,856 essential genes identified based on large-scale CRISPR experiments (Wang *et al*., 2015) were selected to condense the expression information for each cell line. Pearson’s correlation for every pair of cell lines was calculated using the expression fold-changes of these essential genes. Thus, in total, we had 491 and 983 cell line features for CCLE and GDSC, respectively.

For training the model, a drug sensitivity score *s* = −log (*IC*_50_) was defined where the higher the score the more sensitive the cell line is to the drug. Models were trained and tested independently for CCLE and GDSC to avoid biases towards either of the datasets (Haibe-Kains *et al.*, 2013; Haverty *et al.,* 2016).

To train CaDRReS, we used matrix factorization to learn a ‘pharmacogenomic space’ i.e. a latent space to project drug and cell line data such that the dot product between a cell line vector and a drug vector provides the cell-line specific drug response (**Figure 1A**). Drug sensitivity models were then computed based on equation 1:

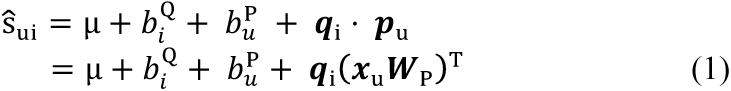

**Fig. 1.**
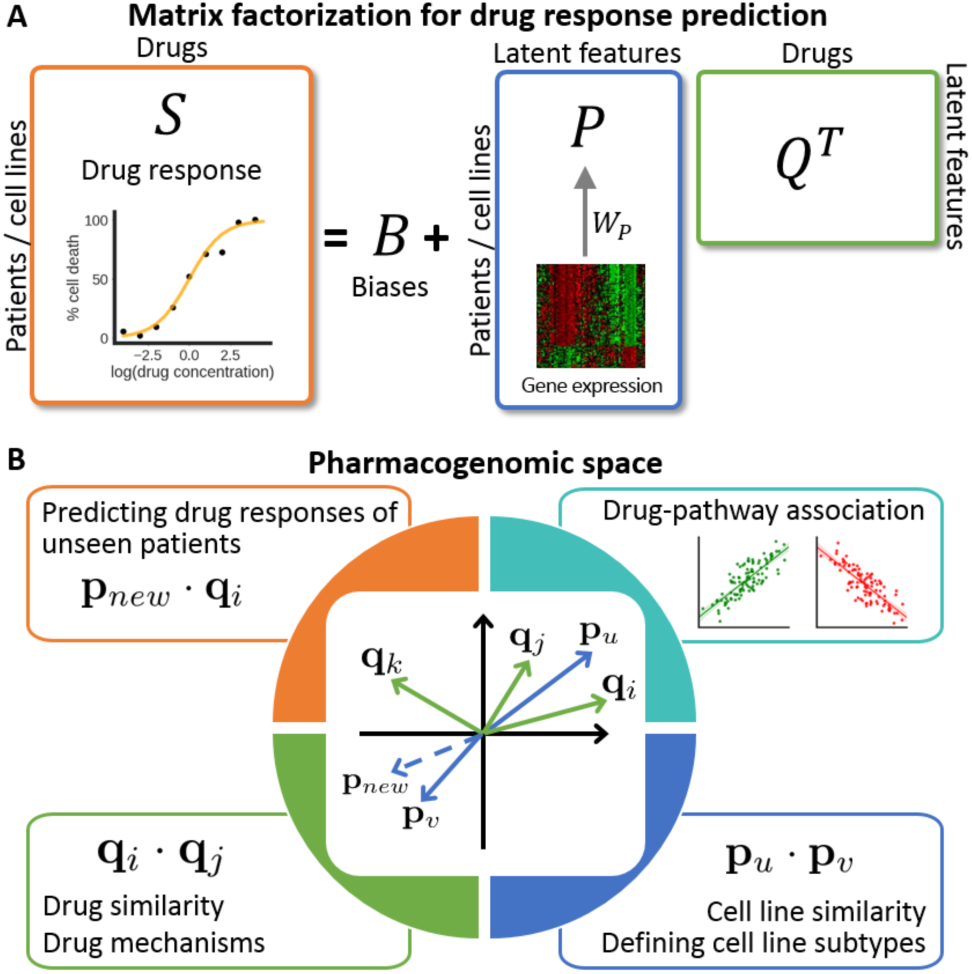
Overview of the CaDRReS framework. Schematic depicting the relationship between the drug response matrix *S*, the bias terms and factorized matrices for cell lines and drugs. A transformation matrix (*W*_*P*_) is used for projecting cell lines onto the latent space. (B) The pharmacogenomic latent space captures interactions between drugs and cell lines and thus enables the study of drug-pathway associations, drug mechanism similarity, and cell line sub-types as discussed in later sections.

where 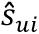 is the predicted sensitivity score of cell line *u* to drug *i*, μ is the overall mean drug response,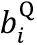 and 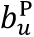 are bias terms for drug *i* and cell line *u*, respectively, ***q***_*i*_,***p***_*u*_ ∈ ℝ ^*f*^ are vectors for drug *i* and cell line *u* in the *f*-dimensional latent space and ***W***_*P*_ ∈ ℝ^*d*×*f*^ is a transformation matrix that projects cell line features ***x***_*u*_ ∈ ℝ^*d*^ onto the latent space. The value of *f* was set at 10 for both CCLE and GDSC datasets based on cross validation performance. As shown in **Figure 1A**, this can be depicted as drug response matrix **(S)** being factorized into biases ***(B)*** and matrices of cell lines ***(P)*** and drugs (*Q*). Rows of the cell line matrix (***P***) and the drug matrix (***Q***) are vectors of cell lines and drugs in a latent space, respectively. The latent *pharmacogenomic space* captures interactions between drugs and the genomic background of cell lines such that the dot product between a cell line vector and a drug vector (***p* ⋅ *q***) represents the interaction between the drug and the cell line. As shown in **Figure 1B** (center), cell line *u* is sensitive to drug *i* and drug *j* while not being sensitive to drug Similarly, cell line *v* is unlike cell line *u* and does not respond to drugs *i* and *j.* This representation thus has many applications including (i) predicting drug responses of unseen samples (cell lines or patients), (ii) revealing drug mechanisms and (iii) subtypes of cell lines, and (iv) identifying drug-pathway associations (**Figure 1B**) as will be discussed in later sections.

In order to train the model the following ‘sum of squared error’ loss function was optimized:

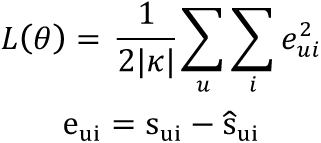

where *s*_*ui*_ and 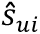 are observed and predicted sensitivity scores for cell line *u* using drug *i,* respectively, ***θ*** = {*b*_*i*_, *b*_*u*_, ***W***_*P*_, ***q***_*i*_}, and *|κ|* is the number of drug response experiments in the training dataset. Finally, we applied gradient descent to optimize this loss function and obtain all parameters in ***θ*** (see **Supplementary method 2**). We tested CaDRReS’ robustness by constructing 10 different models from different random starting points for the gradient descent optimization and observed that the models show similar performance (**Supplementary figure 3**).

### 2.3 Comparisons with Related Methods

We compared the predictive performance and robustness of CaDRReS against other existing methods including a method based on the elastic net regression model (**ElasticNet**; Barretina *et al.,* 2012; Iorio *et al.,* 2016), **cwKBMF** (Khan *et al.,* 2016), the method from **Sheng et al** (Sheng *et al.,* 2015), as well as a control method based on random permutations of the drug sensitivity scores for each cell line (**Control**). For ElasticNet, the model was trained for each drug as described previously (Barretina *et al.,* 2012; Iorio *et al.,* 2016) using the Elastic Net library from Scikit-learn (*l*1*-* ratio = 0.5; Pedregosa *et al.,* 2011), where the model automatically selects the genes. For the method proposed by Sheng et al (Sheng *et al.,* 2015), we re-implemented it as described in the paper, normalized drug response data, calculated drug similarity and drug-specific cell line similarity scores and set the parameters *r_d_* (number of similar drugs) = 3 and *r_c_* (number of similar cell lines) = 9 as used in the paper. For cwKBMF, drug response data was normalized for each drug as described in the paper and the provided MATLAB source code was used to train a model.

### 2.4 Evaluation Metrics

We performed 5-fold cross-validation to evaluate the predictive performance of the models. For evaluating cell line ranking for each drug, we calculated Spearman correlation (*r*_*s*_) and reported the average correlation across drugs. To evaluate models for each cell line, the normalized discounted cumulative gain (NDCG), a widely used score for evaluating ranking recommendations, was calculated as follows:

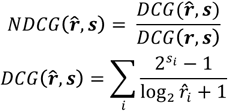

where 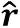 is the predicted rank of drugs tested on a cell line, ***s*** is a list of observed drug sensitivity scores and ***r*** is the known ranking of drugs. NDCG ranges from 0 to 1, where 1 indicates that the model correctly predicts the ranking of drugs. The numerator in DCG is designed to give greater weight to a drug with higher sensitivity score, while the denominator gives preference to drugs predicted to have higher ranks.

### 2.5 Identifying Drug-Pathway Associations

Using 217 Biocarta pathway gene sets from MSigDB (Liberzon et al., 2011), pathway activity scores were calculated for each cell line by summing up gene expression fold-changes of genes in each pathway. To identify drug-pathway associations, we then calculated the Pearson correlation between pathway activity scores and predicted drug responses (**log (*IC*_50_**); lower values indicate greater response), where a negative correlation suggests that a pathway is essential for drug effectiveness, while a positive correlation suggests that it plays a role in drug resistance.

## 3 Results

### 3.1 Performance and Robustness of CaDRReS

A common way to evaluate drug response prediction methods is to assess their correlation (or squared error) compared to known responses for each drug (across cell lines) in a cross-validation framework (Barretina *et al.,* 2012; Iorio *et al.,* 2016). Using the matrix-factorization based approaches, CaDRReS and cwKBMF showed significantly better performance than ElasticNet, Sheng et al, as well as the Control method *(p-*values <10^-30^) in both the CCLE and GDSC datasets (**Figure 2A**). While the ability to predict cell line responses for a given drug is useful to understand drug efficacy and to characterize drug mechanisms, ranking drugs for a given unseen cell-line/patient may be more relevant for precision oncology applications. Based on a weighted scoring of rankings (NDCG), we noted that CaDRReS and ElasticNet exhibited similar performance and improved notably over cwKBMF, Sheng et al, and the Control method (*p*-values **<10^-20^; Figure 2B).** Taken together, these results suggest that CaDRReS improves over existing approaches in providing models that are useful for both drug response prediction across cell-lines and within a cell-line.

**Fig. 2.**
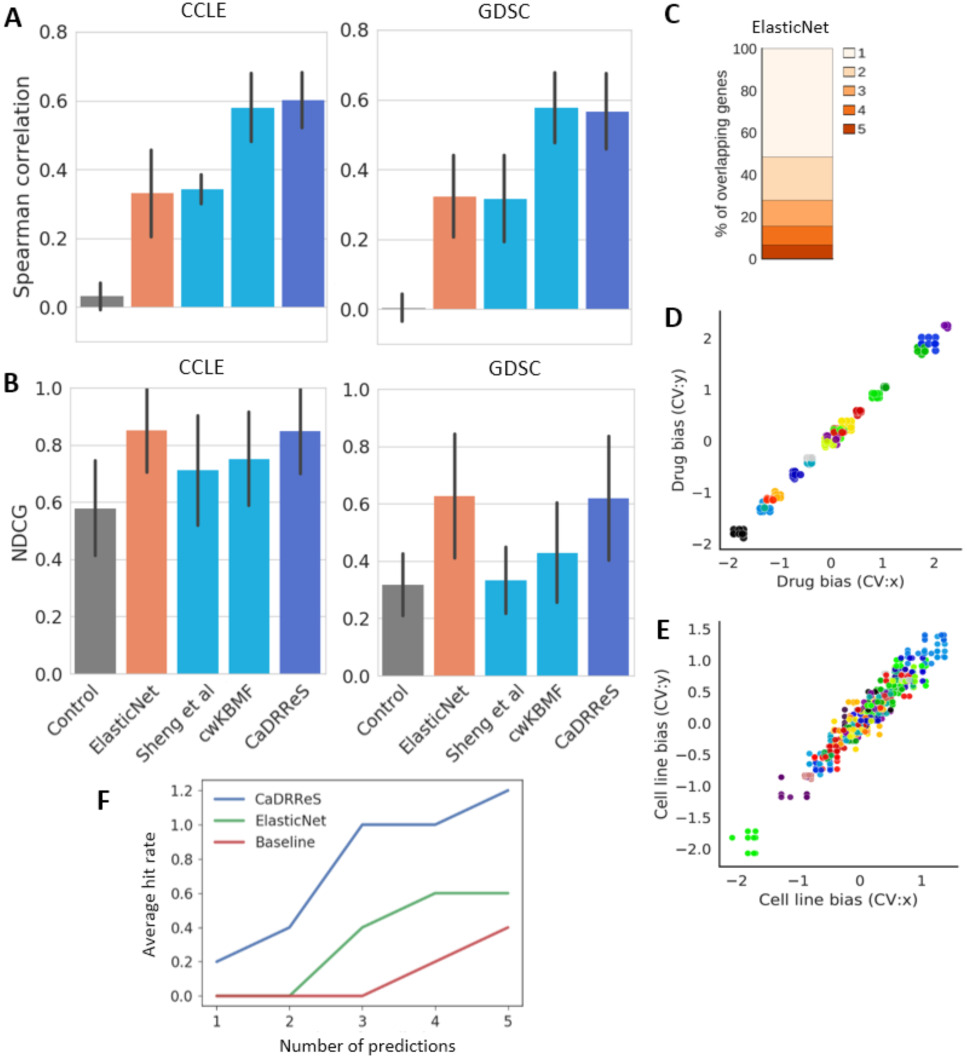
Performance and robustness of the CaDRReS model. (A) Average performance (spearman correlation) across drugs based on 5-fold cross-validation (error bars represent 1 standard deviation). (B) Average NDCG scores across unseen cell-lines based on 5-fold cross-validation. (C) Average percentage of overlapping genes in ElasticNet across different CCLE cross-validation datasets. (D) Concordance between drug-specific bias terms as inferred by CaDRReS for every pair of cross-validation runs. Each color represents a drug in the CCLE dataset. (E) Concordance between cell line bias terms as inferred by CaDRReS for every pair of cross-validation runs. Each color represents a cell line in the CCLE dataset (first 50 cell lines). (F) Average hit rate (number of sensitive drugs identified) in the top five predictions of each method. Baseline refers to an approach that sorts drugs by their average sensitivity across cell lines.

For drug response prediction within a cell-line, although ElasticNet models were trained independently for each drug, their NDCG scores were surprisingly high. We suspected that this may be due to overfitting while training using a limited number of cell lines for each drug. To assess this, we evaluated the robustness of ElasticNet models learned across cross-validation runs and found that <10% of the selected genes were shared across folds and half of the genes were selected in only one fold **(Figure 2C).** In contrast, CaDRReS showed consistently high correlation for drug biases (0.99; **Figure 2D**) and cosine similarity of inferred drug vectors (0.96) across cross-validation runs, as well as high correlation for cell line biases (0.96; **Figure 2E**) and cosine similarity of the inferred cell line vectors (0.88), highlighting the robustness of its models.

To further evaluate their performance, CaDRReS and ElasticNet models were trained on the GDSC dataset and tested on an independent dataset from patient-derived HNC cell-lines. Sheng et al and cwKBMF were not included here because they require normalization of drug response data, which leads to a loss of drug ranking information within a cell line. Despite having similar performance on the GDSC dataset, CaDRReS outperformed ElasticNet on this independent dataset **(Figure 2F),** emphasizing its ability to provide more robust and generalizable models. In particular, CaDRReS was able to identify on average at least one drug that elicited a strong response for each cell-line among its top 3 predictions, while a baseline method based on average response across cell lines identified none.

### 3.2 Investigating Drug Mechanisms via the Pharmacogenomic Space

We trained CaDRReS models on the full datasets to obtain drug and cell-line biases, as well as the pharmacogenomic spaces capturing drug-drug, cell line-cell line, and drug-cell line associations for both CCLE and GDSC **(Supplementary figure 4).** Then to study drug mechanisms, we took vectors defined for each drug in the pharmacogenomic space, computed cosine similarities between every pair, and compared these to a commonly used drug structural similarity score (Tanimoto coefficient of SMILES calculated using the SMSD toolkit; Rahman *et al.,* 2009). Drug cosine similarities were significantly higher for drug pairs having high structural similarities (Tanimoto coefficient > 0.3; Wilcoxon test *p*-value <0.04 for CCLE and <0.001 for GDSC), suggesting that in general, similarly structured drug pairs tend to have higher cosine similarity on the pharmacogenomic space and thus elicit similar responses **(Supplementary figure 5).** However, there are indeed exceptions to this rule where drugs that elicit similar response profile have significantly different chemical structures. For instance, PD-0332991 and PHA-665752 have relatively low structural similarity (Tanimoto coefficient = 0.07), but high correlation of the observed drug responses (0.51 with *p-*value < 10^-29^). This is likely due to the fact that PD-0332991 is a CDK4/6 inhibitor that can reduce RB phosphorylation (Fry *et al.,* 2004), while PHA-665752 can inhibit c-MET and thus result in reduced phosphorylation of RB down-stream (Ma *et al.,* 2007). Thus drug similarity in the pharmacogenomic space has the potential to capture deeper similarities in drug response mechanisms beyond those observed purely based on drug structural similarity.

In the pharmacogenomic space, we observed that clusters of drugs frequently represent groups that target the same gene or pathway **(Figure 3A, Supplementary figure 6).** For example, EGFR inhibitors (Lapatinib, ZD-6474, AZD0530, Erlotinib), RAF inhibitors (RAF265, PLX4720) and MEK inhibitors (PD-0325901, AZD6244) in CCLE formed separate clusters based on cosine similarity. In addition, cosine similarities among the five MEK1 inhibitors in GDSC (CI-1040, PD-0325901, RDEA119, Trametinib, and selumetinib) were significantly higher than between MEK1 inhibitors and other drugs (p-value <10^-15^). A similar trend was also observed for the four BRAF inhibitors, AZ628, Dabrafenib, PLX4720, and SB590885 *(p-*value <10^-7^; **Figure 3B**). These observations are interesting given that CaDRReS was trained based solely on drug response data, without any other information on drug properties.

**Fig. 3.**
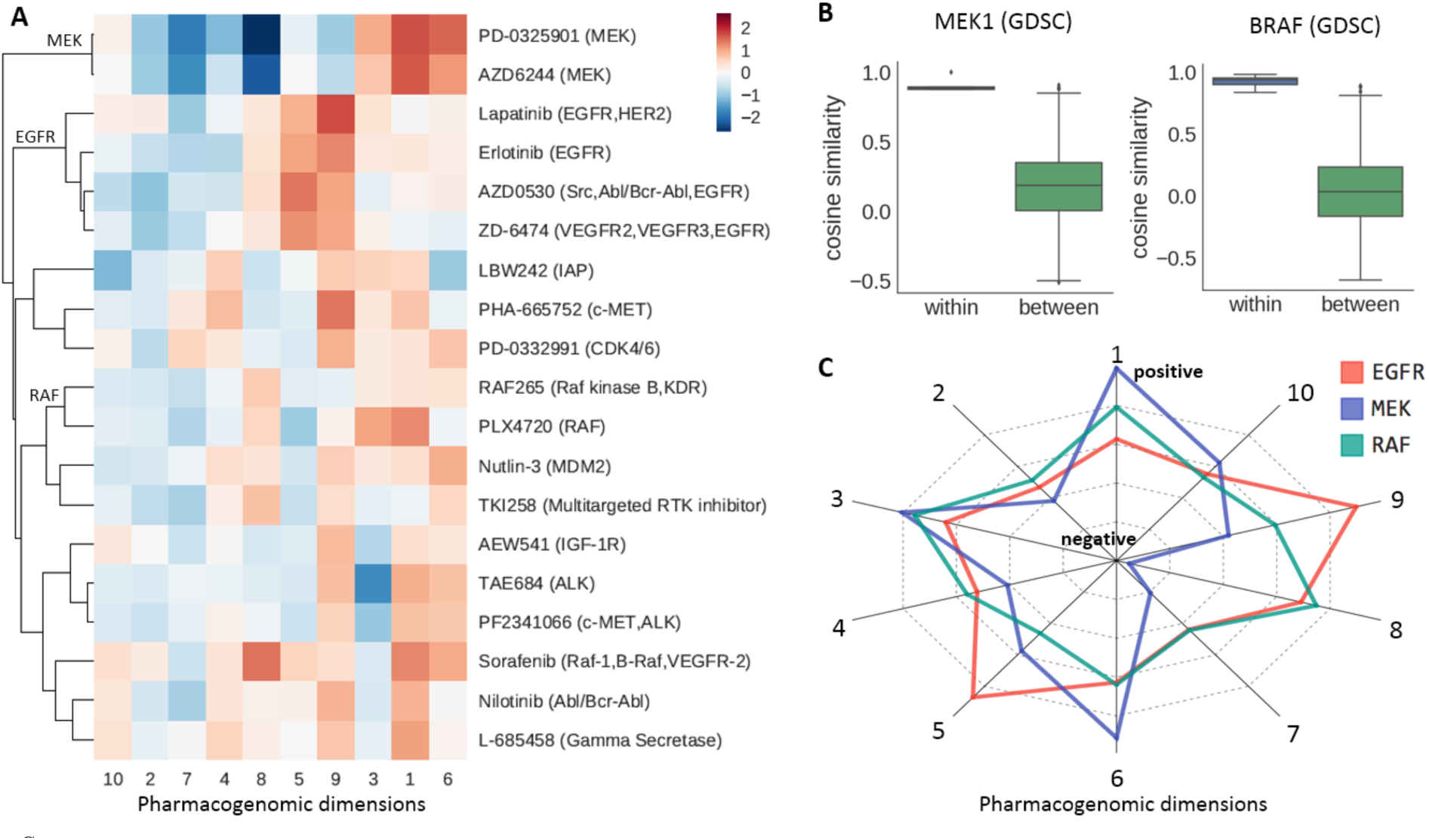
Clustering of drugs on the pharmacogenomic space and its relation to mechanism-of-action. (A) Heatmap presenting average linkage hierarchical clustering of drugs based on cosine similarity on the pharmacogenomic space (CCLE). (B) Distribution of within-and between-group cosine similarities of drugs targeting MEK1 (GDSC) and BRAF (GDSC). (C) Representation of dimensions of the pharmacogenomic space capturing different drug mechanisms. For each target, the average vector of the corresponding drugs was calculated for EGFR, RAF, and MEK inhibitors (CCLE).

By examining dimensions of the pharmacogenomic space, we observed that each dimension captured different aspects of sensitivity to various drug classes **(Figure 3C).** For example, EGFR inhibitors dominated in the 5th and 9th dimensions and thus cell lines that were projected close to the positive sides of these dimensions have higher EGFR inhibitor sensitivity. Additionally, we observed that MEK inhibitors lie on the negative side of the 8^th^ dimension and the values of cell line vectors in this dimension were most positively correlated with activity scores for the EIF2 pathway (0.217), indicating that cell-lines with inactivated EIF2 pathway may be more sensitive to MEK inhibitors. This observation is in agreement with prior work showing that MEK inhibitors work by inducing activation of eIF-2B, which results in a shutdown of cellular protein synthesis and leads to apoptosis (Quevedo et al., 2000; Liberzon et al., 2011). These results highlight the utility of the pharmacogenomic space learned by CaDRReS for capturing interpretable information related to drug mechanisms and pathways.

### 3.3 Cell Line Subtypes in the Pharmacogenomic Space

Clusters of cell-lines in the pharmacogenomic space should in-principle be tuned to capture drug-response similarities. However, not surprisingly we found that they also capture tissue type signatures, with cell-lines from the same tissue type showing significantly higher cosine similarity than cell-lines from different tissue types (**Figure 4A**, **Supplementary figure 7A**), and also being visually distinct in t-SNE (Maaten and Hinton, 2008) 2D space (**Figure 4B**, **Supplementary figure 7B**). Further segregation into histological subtypes was not always as clear (**Supplementary figure 7C**), though most small cell lung carcinoma (SCLC) cell-lines were distinct from non-small cell lung carcinoma (NSCLC) cell-lines (except for NSCLC carcinoid cell-lines; **Figure 4C**). The placement of NSCLC carcinoid cell-lines with SCLC cell-lines is clearly reflected in their drug-response profiles: e.g. while NSCLC cell-lines were typically sensitive to PD-0325901 (MEK inhibitor), carcinoid cell-lines were not (**Supplementary figure 8**). In addition, we found that cell-lines with KRAS mutations had significantly higher predicted PD-0325901 sensitivity (adjusted *p-*value <1.4x10^-8^), and that KRAS mutations were common in NSCLC cell lines (~30%) but not seen often in SCLC or carcinoid cell-lines (~3%), in agreement with prior work on KRAS mutations being activation biomarkers for MEK inhibitors (Stinchcombe and Johnson, 2014).

**Fig. 4.**
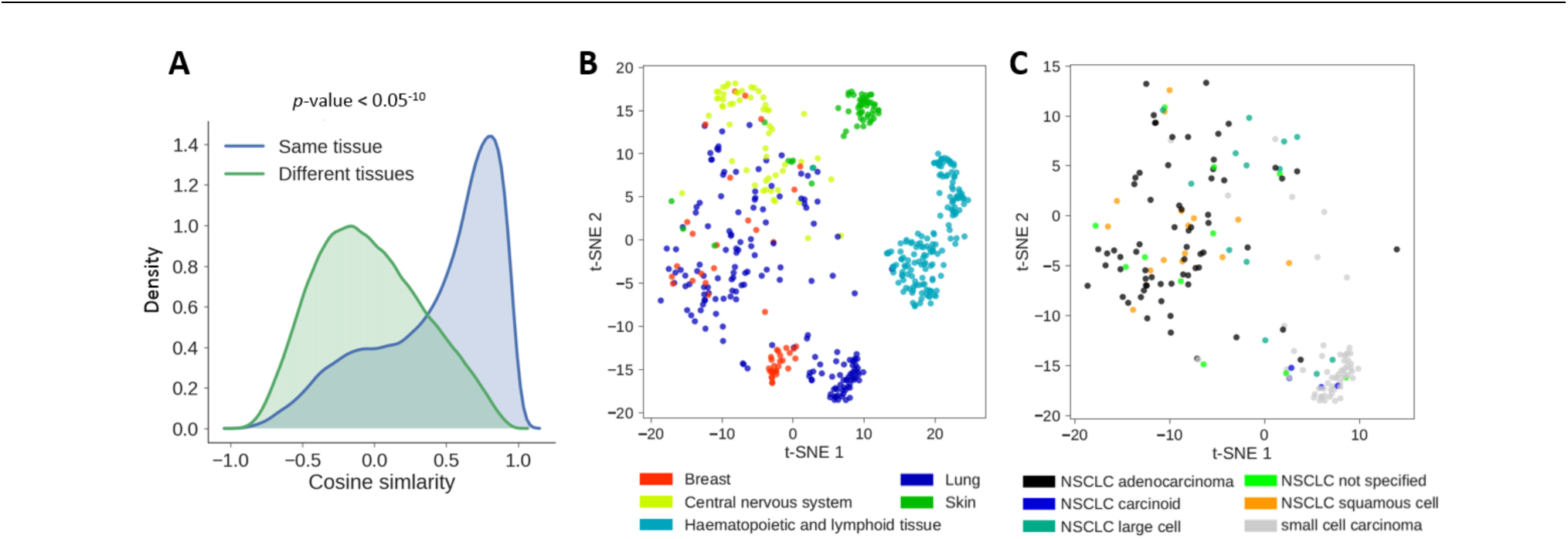
Subtypes of cell-lines on the pharmacogenomic space. (A) Kernel density plot showing distributions of cosine similarities between cell-lines of the same tissue type and of different tissue types (GDSC). (B) Visualization of GDSC cell-lines from top 5 most frequent tissue types using t-SNE. (C) Visualization of different subtypes of GDSC lung cancer cell lines using t-SNE.

By leveraging pathway information, we observed that activity scores for the ERK pathway in NSCLC cell-lines (mean=1.52) were significantly higher than for SCLC cell-lines (mean=-3.24; p-value <1.3x10^-9^), and the activation of ERK pathway due to KRAS mutation could play a role in the increased sensitivity to MEK inhibitors (RAF-MEK-ERK pathway; Stinchcombe and Johnson, 2014). In contrast, cell-lines with RB1 mutations had a significantly lower PD-0325901 sensitivity (adjusted *p*-value <7x10^-8^), and correspondingly RB1 mutations were more common in SCLC cell-lines (67%) than in NSCLC cell-lines (10%). These observations corroborate earlier work suggesting that mutations in the RB1 pathway can inhibit the RAF-MEK-ERK pathway and thus induce resistance to MEK inhibitors (El-Naggar *et al.,* 2009). Cell-line clusters determined by CaDRReS thus correlated well with mutation and pathway activation in explaining drug responses, and could serve to construct new testable hypotheses when such information is not known.

### 3.4 Associations between Drugs and Pathways

Associations between cancer drugs and key pathways can be identified in the pharmacogenomic space based on pathway activity scores, cell line vectors, and drug vectors (see **Methods** and **Supplementary tables 3**, **4**). As expected, we observed that drugs targeting the same gene were frequently associated with the same set of pathways (**Figure 5A**). For instance, four EGFR inhibitors had IC50 values that were negatively correlated with activation scores for the EGFR SMRTE pathway (assistant association), consistent with a study showing that amplification of the EGFR gene is correlated with high response to anti-EGFR agents. (Normanno *et al.,* 2006). Similarly, two RAF inhibitors showed assistant associations with the VEGF-Hypoxia-Angiogenesis pathway (VEGF), in agreement with previous studies showing that VEGF expression induced by Raf promotes angiogenesis, while RAF inhibitors can block the RAF/MEK/ERK pathway and inhibit tumor angiogenesis (McCubrey *et al.,* 2007; Liu *et al.,* 2006).

**Fig. 5.**
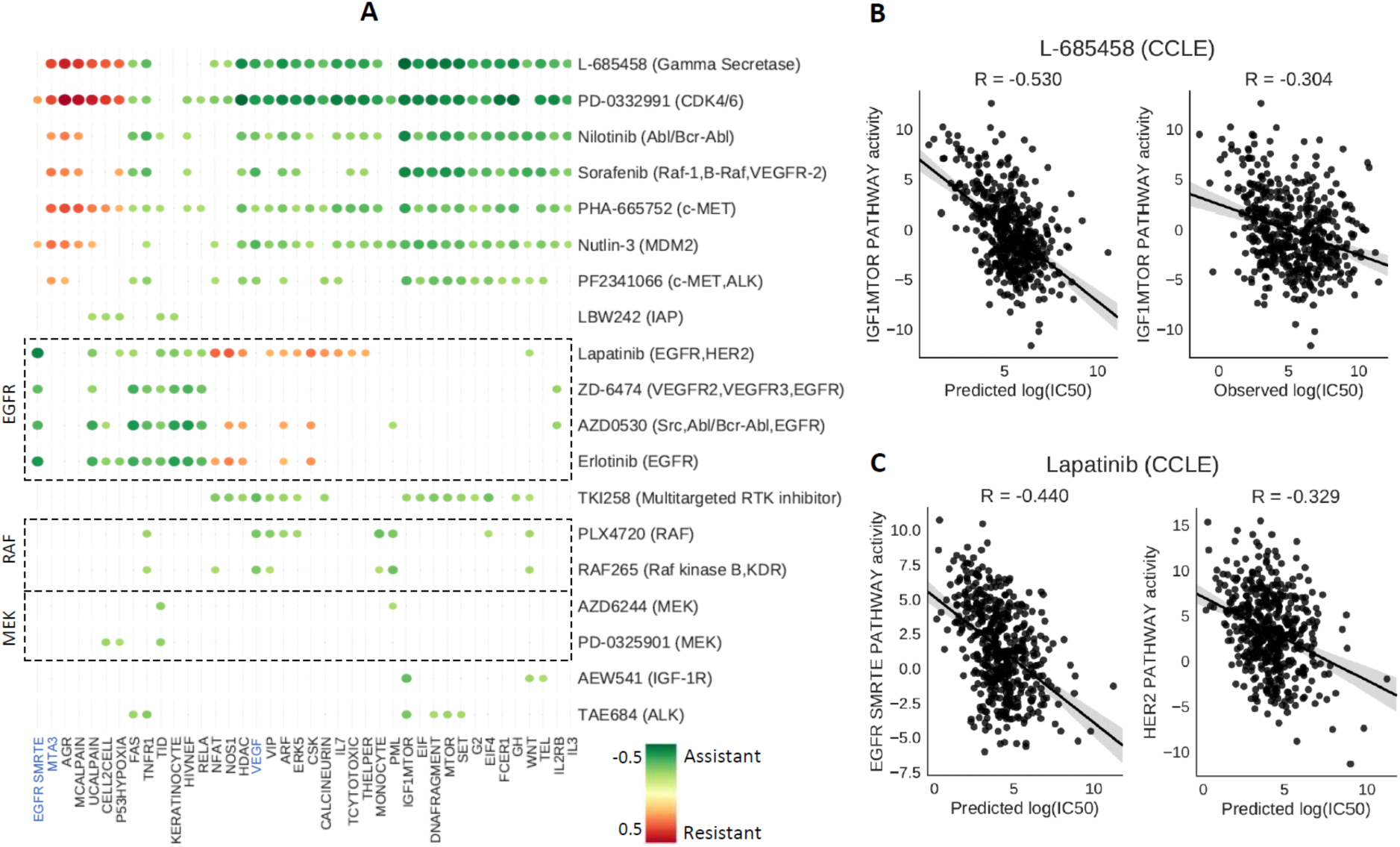
Drug-pathway associations identified on the pharmacogenomic space. (A) Drug-pathway associations based on CCLE data. For visualization, the top 40 pathways having highest associations across drugs (average absolute correlation) were selected. Negative and positive correlations between pathway activity and drug sensitivity scores are denoted as being “assistant” and “resistant” associations, respectively. (B) Assistant associations between L-685458 (gamma-secretase inhibitor) and IGF-1 MTOR pathway. (C) Assistant associations between Lapatinib (EGFR inhibitor) and EGFR SMRTE and HER2 pathways.

We also observed resistant associations between the MTA3 pathway (MTA3) and multiple drugs such as L-685458 (gamma-secretase inhibitor) and PD-0332991 (CDK4/6 inhibitor), suggesting that the cell lines with inactivated MTA3 pathway tend to be sensitive to these drugs. In addition, the study of Fujita *et al.* showed that the absence of MTA3 leads to invasive growth in breast cancer (Fujita *et al*., 2003). Taken together, these observations suggest that drugs having resistant association with MTA3 pathway might be effective when tumor growth is caused by the downregulation of the MTA3 pathway, although further work is needed to confirm this hypothesis.

In terms of drug-pathway associations, we noted that the strongest assistant association was observed between the drug L-685458 (gamma-secretase inhibitor) and the IGF-1 MTOR pathway (**Figure 5B**). This observation is also borne out in studies reporting that gamma-secretase inhibitors can inactivate MTOR signaling pathway and consequently induce apoptosis (Shih and Wang, 2007). Interestingly, we observed a stronger association signal for predicted drug responses than observed drug responses, suggesting that CaDRReS may have an ability to reduce the noise observed in experimental drug response data. Stronger signals based on predicted drug responses were also observed for other known assistant associations, such as the one between Lapatinib (an EGFR inhibitor) and the EGFR SMRTE pathway (R=−0.440 vs −0.329) as well as the HER2 pathway (R=-0.288 vs −0.242) (**Figure 5C;** Harari, 2004; Medina and Goodin, 2008). These results highlight the utility of predictions from CaDDReS for discovering pathway biomarkers for drug sensitivity.

## Discussion

Several drug response prediction models have been proposed in the literature, with a primary focus on predicting the response of different cell-lines to a given drug. Correspondingly, the performance of these models was evaluated for each drug individually based on the correlation between the predicted and observed drug responses. However, while predicting cell-line response to each drug may provide insights into differential drug response mechanisms, the ability to rank drugs for unseen cell-lines/patients is likely to be more useful from a clinical perspective. Therefore in this work, besides evaluating response correlations for each drug, we evaluated the ability to correctly order drugs for a given cell-line using a popular weighted metric for rankings (NDCG). Under both these metrics of evaluation, CaDRReS consistently provided the best models and was also able to perform well on unseen datasets.

In addition to its robust models, a useful feature of CaDRReS is the ease with which its models can be interpreted, an aspect that has not been given the attention it deserves in earlier studies. Models trained by CaDRReS provide a projection of cell-lines and drugs into a *pharmacogenomic space* which can be used to explore drug-drug, cell line-cell line, and drug-cell line relationships as shown in sections 3.2-3.4. This is in addition to the easy visualization and clustering analysis that this representation permits (e.g. **Figure 3** or **Figure 4**). In contrast, while the ElasticNet model provides high concordance between observed and predicted cell line rankings, non-robustness in gene selection means that it may not be meaningful to biologically interpret the selected set of genes for a given drug. Similarly, while the cwKBMF model incorporates pathway information and can be used to infer the strength of drug-pathway associations, it does not provide directionality for these associations. CaDRReS models start off by being agnostic of pathways but by incorporating this information later, allow us to identify both strength and directionality of drug-pathway associations as highlighted in the results in **Figure 5**.

Currently, drug response prediction models are trained on drug response data for cancer cell lines, but ignore the toxicity of drugs due to the unavailability of corresponding information using normal cells. This likely limits the practical utility of such models as drugs that elicit a strong response across cell lines may also have higher toxicity. Refined models that take into account drug toxicity could also find application in studying drug synergies using the pharmacogenomic space: the sum of drug vectors could be used to predict synergistic response, and thus enable the goal of reducing drug dosage to limit side-effects.

An important limitation for the field of drug sensitivity prediction is that despite the presence of several publicly available cancer drug-screening datasets, the number of cell types and drugs in each dataset is still limited compared to the complexity of the models. Being able to merge information across multiple datasets could thus help construct more robust and general models. Experimental inconsistencies and noise across datasets have so far, however, stymied efforts to work towards this goal (Haibe-Kains *et al.,* 2013; Haverty *et al.,* 2016).

Although CaDRReS was among the top performing models for both cell-line and drug ordering in **Figure 2**, it still considered only gene expression of essential genes in its models. We suspect that integrating other types of omics data, such as mutations, in a meaningful manner can enrich information in the dataset and thus improve the predictive performance of corresponding models. Additionally, using information from gene interaction networks to capture relationships between genes could be another way to improve the performance and interpretability of this model in the future.

## Acknowledgements

We thank Dr. Lie Yong Judice Koh for providing drug response data for patient-derived HNC cell lines and suggestions on the manuscript. We also thank Dr. Jonathan Goke for valuable insights and comments on the manuscript.

